# Negative density-dependent parasitism in a group-living carnivore

**DOI:** 10.1101/2020.06.15.153726

**Authors:** Gregory F Albery, Chris Newman, Julius Bright Ross, Shweta Bansal, Christina Buesching

## Abstract

Animals living at high population densities commonly experience greater exposure to disease, leading to increased parasite burdens. However, social animals can benefit immunologically and hygienically from cooperation, and individuals may alter their socio-spatial behaviour in response to infection, both of which could counteract density-related increases in exposure. Consequently, the costs and benefits of sociality for disease are often uncertain. Here, we use a long-term study of a wild European badger population (*Meles meles*) to investigate how within-population variation in host density determines infection with multiple parasites. Four out of five parasite taxa exhibited consistent spatial hotspots of infection, which peaked among badgers living in areas of low local population density. Combined movement, survival, spatial, and social network analyses revealed that parasite avoidance was the likely cause of this negative density dependence, with possible roles for localised mortality, encounter-dilution effects, and micronutrient-enhanced immunity. These findings demonstrate that animals can organise their societies in space to minimise parasite infection, with important implications for badger behavioural ecology and for the control of badger-associated diseases.

## Introduction

A wild animal’s infectious disease burden is determined by its exposure and susceptibility to infective pathogens. Typically, higher population density results in increased contact rates and thus greater exposure [1,2]; however, sociality carries immunological and hygienic benefits that can counteract this exposure-exacerbating effect [3,4]. For instance, cooperative foraging improves nutrition, alleviating infection costs [5]; mutual grooming removes ectoparasites [6–8]; and “social immune responses” maintain collective group health [9,10], e.g. through extirpating sick individuals [11]. Additionally, the environmental distribution of resources can influence spatial behaviour [12–14] and determine local population densities [15], while altering susceptibility [16] and transmission efficiency [17]. Sociality and disease may thus be confounded through shared causal origins rather than being mechanistically linked [17]. Finally, individuals can minimise exposure by avoiding environmental cues [18,19] or infected conspecifics [8,20], creating a population-level “landscape of disgust” analogous to predatory “landscapes of fear” [21,22]. These processes could produce negative density effects, sometimes depending on parasite transmission mode [1,17,23], but their role in defining observed density-infection relationships is poorly understood.

Uncertainty concerning density-infection relationships largely arises because studies of social effects are often carried out across discrete groups [1,24], between populations [25,26], or between species (e.g. [1,2,24,27]). Continuous density measures are rarely linked to parasite burden within one contiguous host population, and studies that do so generally use purely social metrics rather than spatial density gradients (e.g. [28–30]). Continuous spatial density measures are advantageous because: density is a continuous, spatially explicit variable [31–33]; for the many parasites that are spread through the environment, spatial density measures better represent transmission than direct (social network) metrics [17,34–36]; and between-population and cross-species comparisons are fraught with confounding factors such as shared environmental causality or compensatory evolutionary changes [1,3], which within-population studies can more easily avoid or overcome. Furthermore, accounting for spatial effects may help anticipate or test the caveats of cooperation, environmental causality, and behavioural avoidance outlined above.

European badgers (*Meles meles*) are nocturnal, ground-dwelling mustelids with a pan-European distribution, eating a varied diet composed largely of earthworms [37]. Badgers are of particular interest for cattle health in the United Kingdom because of their role in maintaining and spreading bovine tuberculosis [37,38]. Although intended as a control measure, intense culling perturbs badger social systems, causing stressed survivors to disperse, thereby spreading bovine tuberculosis to neighbouring farms [39–41]. Understanding badgers’ socio-spatial behaviours and their epidemiological implications is therefore an important research priority (e.g. [38,42–44]). The demography, behaviour, and parasitism of badgers in Wytham Woods, Oxfordshire have been studied continuously since 1987 [37,45]. Dens (termed “setts”) are situated dependent on soil composition and landscape topography [46]. These badgers reside in cohesive mixed-sex social groups, with around 35% consistent philopatry; however, these groups are also fluid: 16.4% of individuals are trapped at a different social group to their previous capture history, and 19.8% away from their natal group [47]. Furthermore, from genetic pedigree, 48% of cubs have extra-group paternity [48]. Badgers host several arthropod parasites, including badger-specific fleas (*Paraceras melis*) and lice (*Trichodectes melis*) and generalist ticks (*Ixodes* sp.) [49]. They also carry two gastrointestinal protozoans: *Eimeria melis* and *Isospora melis. E. melis* predominantly infects young individuals, causing substantial mortality [50,51]. Although studies have examined social grooming effects on ectoparasite burden [6,52] and roles of denning behaviour in parasite transmission [49], the within-population spatial-social parasite epidemiology has yet to be investigated.

Here, we investigate parasite burdens in the Wytham badger population and their associations with spatial and social behaviour. We establish parasite distributions using spatial autocorrelation models. We quantify social drivers using both spatial density gradients and social network metrics, postulating that greater badger densities would drive higher parasite burdens through increased exposure. Finally, we examine survival effects, investigating whether parasite-linked mortality might alter spatial patterns of badger population density. We consider a range of potential social/spatial covariates of parasite burden, including density-related exposure changes, benefits of cohabitation, condition/susceptibility effects, encounter-dilution effects, and parasite avoidance behaviours.

## Methods

### Data collection

Badgers were sampled as part of a long-term study in Wytham Woods, Oxfordshire, UK (51.778°N, 1.313°W), established in 1987 and recording one of the highest local badger densities ever reported [37,45]. Badgers were trapped overnight at their setts in steel-mesh cages baited with peanuts, collected the next morning, transported to a handling facility and sedated. Individuals were identifiable by tattoo, applied on first trapping (typically as cubs). Measures were taken of head-body length (mm) and weight (to the nearest 0.1kg). We calculated a standardised index of body condition dividing log(weight) by log(body length). The population was trapped seasonally (winter: Jan/Feb/March; spring: April/May/June; summer: July/Aug/Sept; autumn: Oct/Nov/Dec) in 4 quadrants, for 3 days per quadrant. Our dataset included 9016 captures of 1369 badgers spanning 29 years (1990-2018).

The population currently comprises 23 social groups (mean group size=11; range 2-29 individuals), each using more than one sett, with sett/social group affiliation established using baitmarking [46,53]. Badgers were assigned to groups using established residency rules [48]. We computed social networks based on co-trapping using a “gambit of the group” approach, where individuals trapped in the same sett in the same year were assumed to have interacted [54].

Fleas were counted during a stereotyped 20-second inspection of the badger’s full body, and ticks (*Ixodes* sp.) and lice (*Trichodectes melis*) were counted within a 4×4cm square of preferentially-parasitised skin near the groin (per Cox *et al*. 1999). Faecal sampling for two protozoan endoparasites, *Eimeria melis* and *Isospora melis*, was carried out through 1993-1997 and 2009-2017 (N=1287 counts). Endoparasite were counted using salt flotation and microscopy [51,55]. Each count was duplicated, and data were reduced to a binary infected/uninfected status (rather than counts) because of their highly overdispersed distribution.

### Statistical Analysis

#### Covariates

Statistical analysis and data manipulation used R version 3.6.0 [56]. All data and code are available at https://github.com/gfalbery/Badgeworks. Models were constructed using the ‘inla’ package [57,58]. Response variables included counts of fleas and lice (negative binomial distribution) and prevalence of ticks, *Eimeria*, and *Isospora*. Explanatory covariates included: Year (continuous); Month (categorical); Age Category (categorical: cub, yearling, and adult); Sex (male and female); and Body Condition (continuous). Continuous covariates were standardised (mean=0; standard deviation=1). Individual Identity and Year were fitted throughout as categorical random effects.

#### 1. Spatial autocorrelation models

To identify generalised spatial patterns we first fitted models accounting for continuous spatial autocorrelation across the population [33,57,58]. We fitted a base model using only Year, Month, Age Category, and Sex, with individual and annual random effects. We then added an SPDE random effect controlling for spatial autocorrelation in the response variable in space, based on individuals’ point locations. We then compared these models using Deviance Information Criterion (DIC) to establish whether accounting for spatial autocorrelation improved model fit; models within 2ΔDIC were judged competitive.

#### 2. Density models

We then fitted models that replaced the SPDE effects with individual measures of local population density. Models included a range of social/spatial density measures, calculated using various methods. Spatial density metrics represent numbers of individuals in a given space, and therefore tread the line between social and spatial behavioural traits [17]. We computed these by creating space use distribution kernels with the ‘adehabitathr’ package in R [59]. We rasterised the usage distribution, assigning each individual a local density based on the raster value for their map location (Figure 1A-B). Measures included: 1) **Lifetime density**: the density of individuals’ centroids across the study period. 2) **Trapping density**: the density of trapping events, incorporating multiple captures of the same individual per year. 3) **Annual density**: the density of individuals’ centroids in the sampling year, calculated from annual density kernels. Only one spatial density measure was allowed in a given model, as the measures correlate (R>0.5) and co-fitting several measures produced spurious, unexpected correlations in the opposite direction expected from data exploration. We also used two social metrics based on co-trapping patterns. **Degree centrality** was the number of unique badgers with whom each individual was trapped in the same sett per year. **Group size** was the total number of individuals trapped in a given social group per year. Following spatial autocorrelation model procedures, we conducted a model selection procedure for behavioural metrics, using 2ΔDIC as the cutoff, and only including the best-fitting metric. We conducted these models for: the overall dataset (N=9016); adults only (age 2+, fitting age in years as a fixed covariate; N=6159); and juveniles only (N=1639). For these models, we display only the best-fitting metric, and for the overall dataset if it exhibited a significant trend (and a subset-only model otherwise).

**Figure 1:**
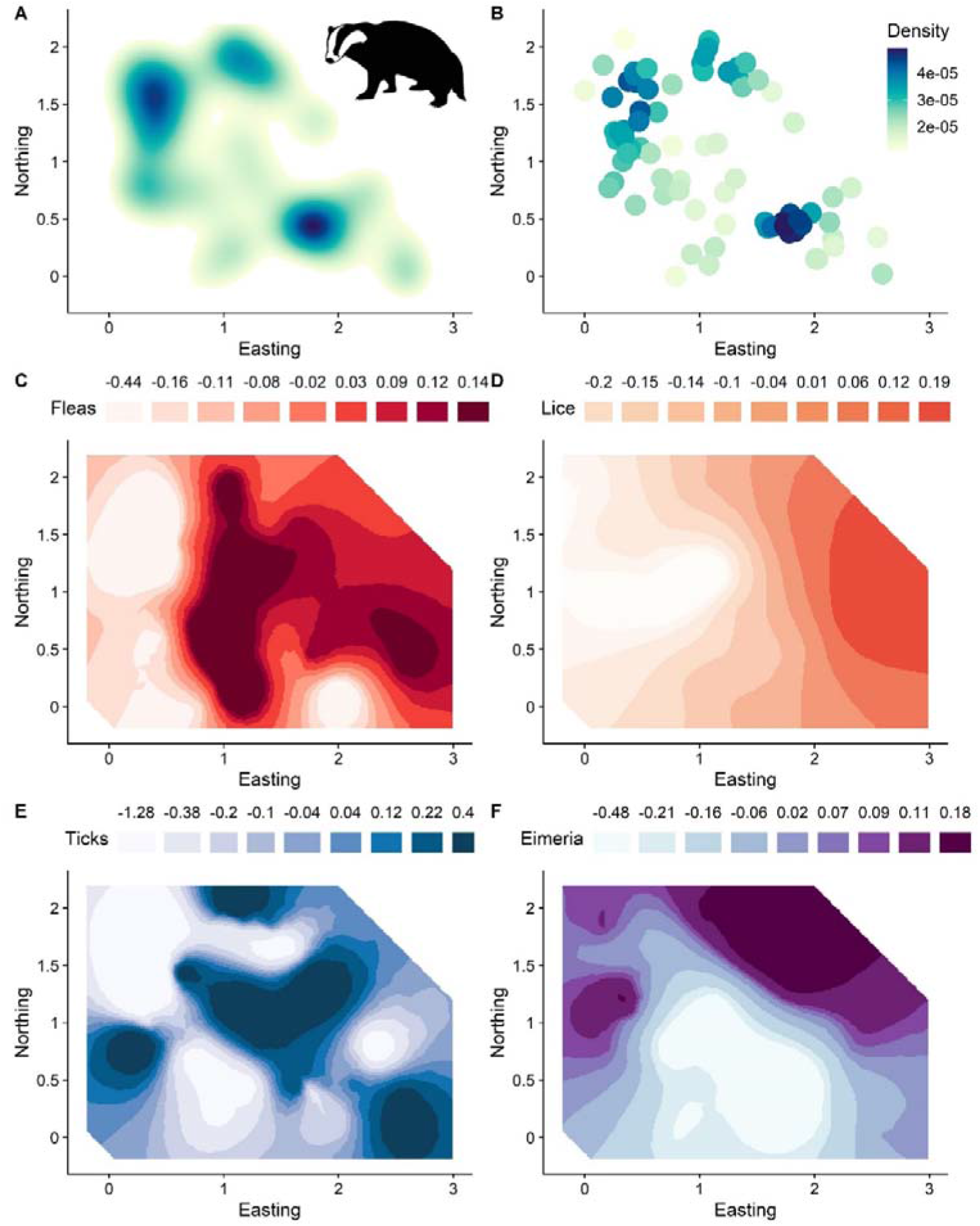
Spatial distributions of badger population density and parasites in Wytham Woods, Oxfordshire, between 1989 and 2018. A: badger population density distributed across Wytham Woods, calculated based on a space use kernel for individuals’ annual centroids. B: Individual badgers trapped at setts (represented by points) were assigned a local density value based on their location on the rasterised space use kernel. Darker blue colours in A and B correspond to greater population density. C-F: the spatial distribution of the four spatially distributed parasites, estimated using the INLA SPDE effect. Darker colours correspond to increased parasitism. The density values in B were fitted as covariates in linear models to explain individual parasite burdens, revealing a negative correlation between density and parasitism (see Fig. 2). All axes are in kilometres, with the 0,0 point at the bottom left of the study area. We examined a fifth parasite, *Isospora melis*, but it was not significantly spatially distributed (see Table 1).

#### 3. Multi-response models

To investigate whether parasites covaried with badger density at the within- or between-individual level, we constructed multi-response models in MCMCglmm with an unstructured covariance matrix [60,61]. We fitted parasites and local density as response variables, estimating their covariance when accounting for other fixed effects and decomposing this covariance at the within- and between-individual level. A negative within-individual (residual) correlation would imply that higher burden/prevalence individuals (compared to their baseline) moved to lower density areas during their lives, supporting either social ostracism or sickness behaviours. In contrast, negative between-individual covariance would imply that individuals inhabiting lower densities generally had inherently greater parasitism. We only constructed these models for parasite-host age category combinations that demonstrated density effects in the INLA models.

#### 4. Survival models

To investigate parasites’ mortality effects, we fitted models with survival as a binary response (1=animal seen in any subsequent year; 0 otherwise). These models included the same covariates as the parasite models, plus parasite prevalence/count and badger density measures. Parasite measures were either log(x+1)-transformed integers (fleas, lice) or binary (ticks, *Eimeria*, and *Isospora*). We conducted another model selection procedure, adding parasite metrics successively if they improved model fit. This was carried out for adults and juveniles separately.

## Results

INLA autocorrelation models revealed considerable spatial structuring of parasite burdens. Models for 4/5 parasites were substantially improved by incorporating spatial effects (ΔDIC>8; Table 1). Only *Isospora’s* model was not improved by spatial autocorrelation (Table 1).

**Table 1:**
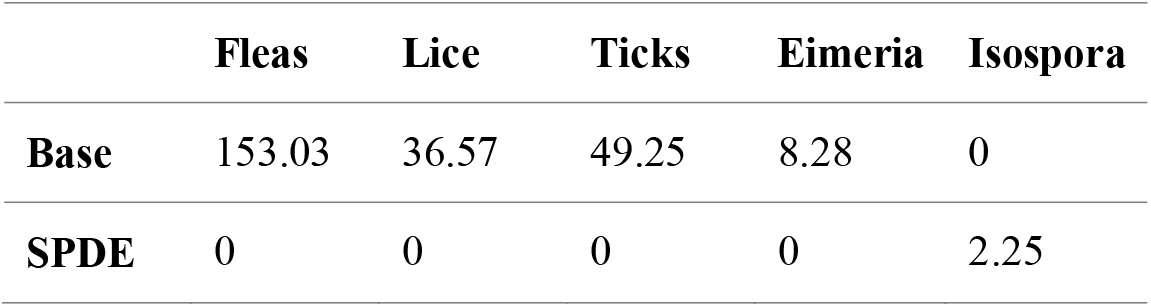
DIC changes associated with spatial autocorrelation terms for the five examined parasites. Lower numbers denote better models; best-fitting models ΔDIC=0. All models except *Isospora* were improved by addition of spatial fields (“SPDE”).

Density models provided substantial support for negative associations between badger density and parasite infection. Including at least one density measure as a covariate improved the model for at least one age category for 4/5 parasites; all had significant negative slopes for density effects (Figure 2; Figure SI1–4; Table SI1–3). Slopes were relatively shallow, but extremely significant and robust (Figure 2). Flea counts were best described by Lifetime Density for the overall model (ΔDIC=-16.31; P<0.0001; Figure 2A), while lice burdens decreased with Trapping Density in the juveniles-only model (ΔDIC=-8.77; P=0.0017; Figure 2B). Tick prevalence was lower in areas of higher Lifetime Density (ΔDIC=-2.47; P=0.0038; Figure 2C), and *Eimeria* prevalence was lower with greater Trapping Density (ΔDIC==-5.83; P=0.0053; Figure 2D). Only one positive social effect was detected: adults with greater degree had slightly more lice (ΔDIC=-2.39; P=0.027; Figure SI2). All DIC changes associated with the model addition procedure are presented in Table SI1–3.

**Figure 2:**
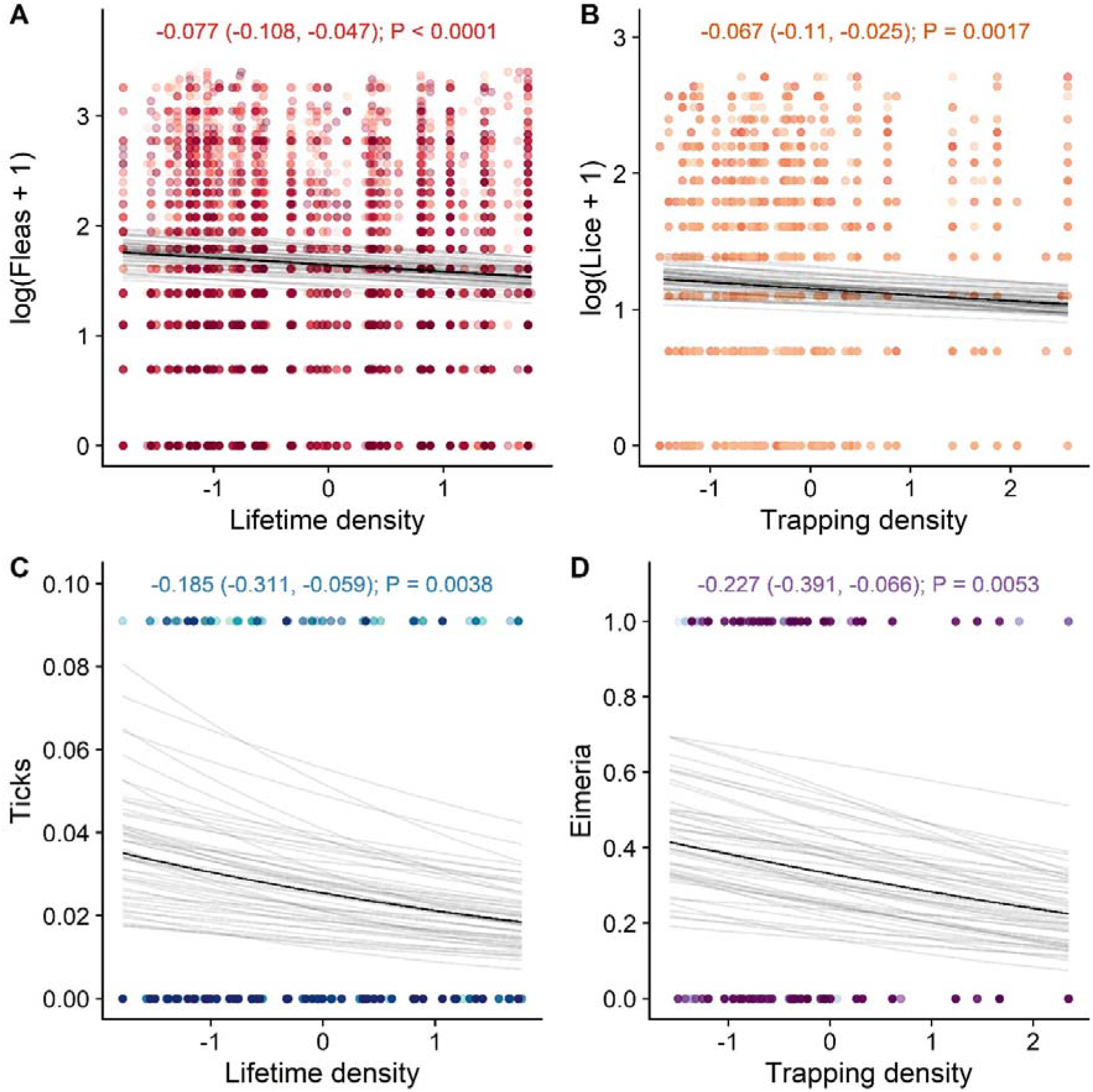
Negative associations between population density and parasite infection. Badgers living at higher densities had fewer fleas (A), fewer lice (B), lower tick prevalence (C), and lower *Eimeria melis* prevalence (D). B represents juveniles only. Points represent individual samples, with colours randomised along a colour palette for plotting clarity. Opaque black lines are derived from linear model fits, taking the mean of the posterior distribution. Transparent grey lines represent 100 fits drawn randomly from the posterior estimate distributions of each model, to demonstrate error in the slope and intercepts. Text at the top of the Figures communicates slope estimates, with lower and upper 95% credibility intervals in brackets, and P values. NB: in panel C, the y axis scale and the location of the positive points have been altered for plotting clarity and to better visualise the slope, due to low tick prevalence.

MCMCglmm multi-response models revealed similar trends to INLA univariate models (Figure SI1–4), and allowed us to decompose density-parasite correlations into within- and between-individual changes (Figure 3B). Correlations were greater and much more significant between-than within-individuals in all cases (Figure 3B). This demonstrates that movement between high- and low-density areas was unlikely to produce changes in parasite burden over an individual’s lifetime, but rather that individuals’ home ranges exhibited inherently different local density-parasite relationships. Only louse infection predicted survival probability; the effect was relatively small, minimally significant, and specific to juveniles (P=0.038; Figure 3A).

**Figure 3:**
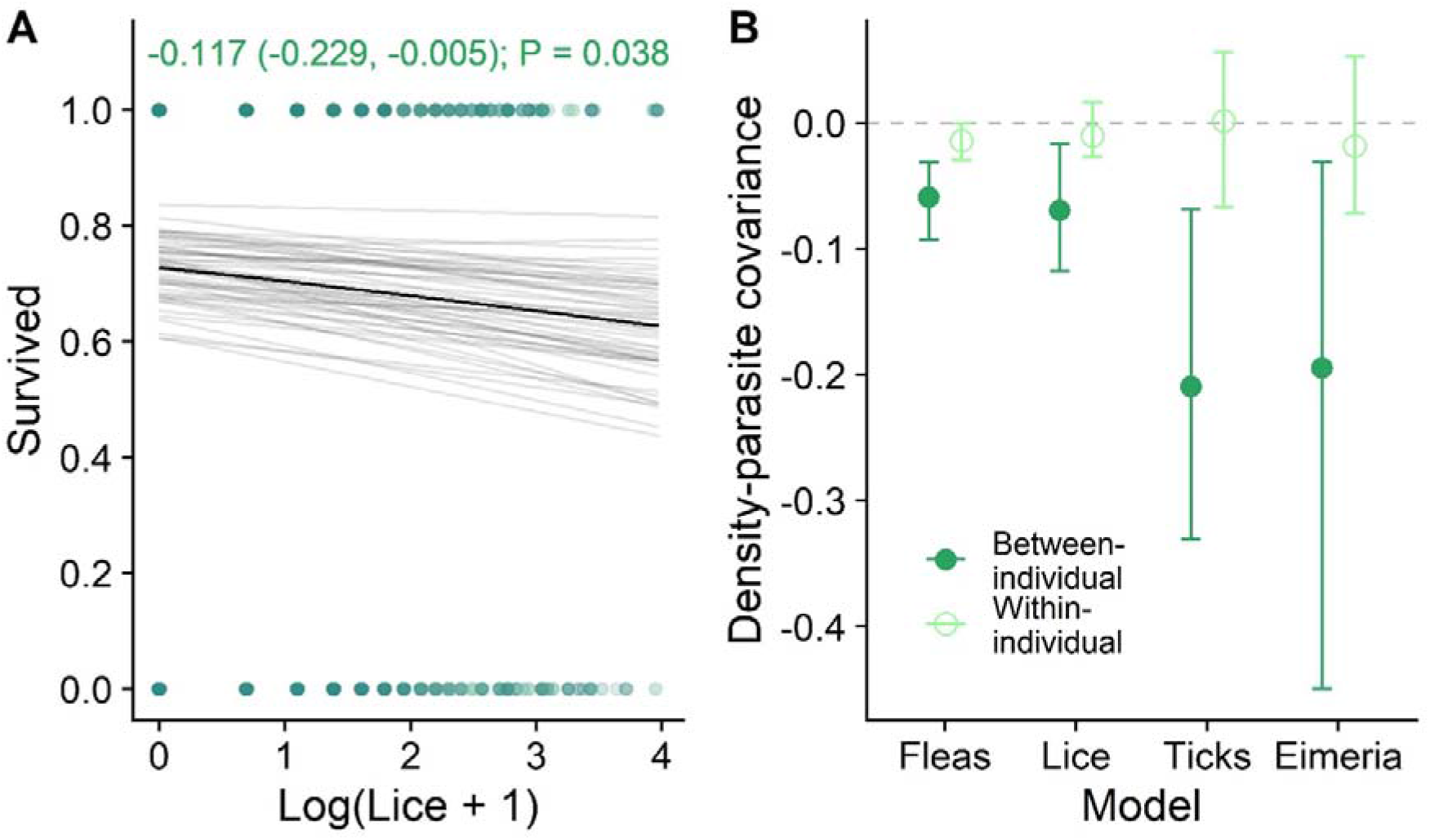
Associations of parasitism with survival and density-parasitism covariance partitioning. A) Louse burden was negatively associated with individual survival probability in juveniles. Opaque black line=linear model fit. Transparent grey lines=100 fits drawn randomly from the posterior estimate distribution, to demonstrate error in the slope and intercept. Text at the top of panel A communicates slope estimate, with lower and upper credibility interval in brackets and P value. B) Estimates for within- and between-individual covariance between parasitism and density, taken from MCMCglmm multivariate models. Points=posterior mean effect size estimates; error bars=95% credibility intervals.

Our models also revealed various other effects (Figures SI1–4). Briefly, in the overall models, juveniles had fewer fleas, more lice, and greater *Eimeria* prevalence than did yearlings and adults, and lower *Isospora* prevalence than adults (Figure SI1). Males had more lice than did females, with substantial monthly variation in all parasites (Figure SI1). Additionally, body condition was negatively associated with fleas, lice, and *Eimeria* infection in all age/sex classes (Figure SI1). The adults-only and juveniles-only models were very similar to the overall models (Figures SI1–3); notably, in adults, lice burden increased with age in years, whereas *Eimeria* and *Isospora* prevalence decreased (Figure SI2).

## Discussion

Using a combination of spatial and social network analysis, we uncovered negative associations between local population density and multiple parasites in this wild group-living carnivore. We found strong but contrasting spatial structuring of fleas, lice, ticks, and *Eimeria*, and, contrary to expectations, all four were most prevalent or abundant in areas of lowest badger density. Co-trapping network and grouping metrics were not predictive of parasitism, implying that “direct” social behaviours, such as mutual allo-grooming, were unlikely to explain the negative density effects. Additionally, badger density effects manifested independent of survival and body condition effects, implying that spatial structuring of the host population did not originate from: 1) localised host mortality, 2) reduced susceptibility arising from co-habitation and implied cooperation benefits [3,5], or 3) greater local resource availability influencing susceptibility [16,62,63]. Additionally, multi-response models revealed low within-individual covariance between density and parasites, providing little support for heavily-parasitised individuals becoming ostracised during the course of their lives [11]. Taking spatial structuring together with setts’ propensity to harbour parasites [49,64], the most parsimonious interpretation is that badgers avoid infection behaviourally, preferring to inhabit areas poorly disposed to parasite transmission (Table 2). These individual-level avoidance behaviours amplify at the population level to produce patterns of badger density inversely related to parasite distributions in the environment, as expected under a “landscape of disgust” [19,21,22]. As well as providing rare evidence of non-consumptive effects of parasites in a wild carnivore, these results imply that animals can arrange their society in space according to parasite transmission.

**Table 2:** Reasoning surrounding our hypothesis testing. We rejected most hypotheses for our observed negative density effects other than parasite avoidance.

| Potential mechanism | Conclusion | Reason for conclusion |
| --- | --- | --- |
| **Co-habitation/**<br>**cooperation benefits** | No | No direct social effects or competition with body condition effects (Model Set 2; Figure SI1-3) |
| **Allo-grooming** | No | Spatial rather than direct social effects; grooming not possible for endoparasites (Model Set 2; Figure SI1-3; Table SI1-3) |
| **Nutrition-associated**<br>**immune benefits** | No | No competition with body condition effects (Model Set 2; Figure SI1-3) |
| **Social ostracism/self-isolation** | No | Low within-individual covariance of density and parasitism (Model Set 3; Figure 3B) |
| **Local host die-offs** | ~No | Possible for lice, in juveniles, but no other mortality effects were evident (Model Set 4; Figure 3A) |
| **Encounter-Dilution** | ~No | Possible for generalist ticks, but not for the other (badger-specific, non-mobile) parasites [77,78] |
| **Avoidance** | Yes | All other possibilities eliminated; consistent with individual-level behavioural responses [64] |

Our study demonstrates observationally that parasites can determine a society’s structure in the environment, and that this phenomenon may counteract the more conventional prediction that host density exacerbates parasitism through increased exposure (e.g. [1,2,65]). Consequently, studies aiming to quantify social covariates of disease should explicitly seek to investigate individual movement and avoidance behaviours, socio-spatial structuring, and feedbacks between sociality and space use [17]. Previous studies on other mammals have revealed avoidance of infected conspecifics [8] and faeces [18,66], but it remained unclear whether animals avoid spatial hotspots of transmission themselves, and whether these could have population-level consequences [21,22]. Badgers respond to social scent cues and may use these to detect and avoid infested or infectious individuals [67]; furthermore, they move between setts regularly to avoid accumulating parasites [64,68], abandoning highly infested setts and chambers [49]; we posit that this behaviour has emergent population-level consequences. Wytham badgers preferentially establish their setts on northwest-facing slopes in areas with sandy soils [46], and variation in internal sett temperature and humidity are associated with reproductive success [69,70]. Our data suggest additional selection for sett traits and sites resistant to parasite infestation and transmission, which produce an emergent trend for fewer setts, with fewer occupants, in more highly parasitised areas. Quantifying parasites in the environment or inside setts and correlating them with badger behaviour could provide support for this hypothesis. Environmental (sett chamber-based) data on parasite presence would be needed to determine whether badgers avoid infection *actively* (pre-infection), using environmental cues that predict parasitism, or *reflexively* (post-infection), by moving away from areas of high parasitism due to irritation or sickness.

Four parasites exhibited negative density effects, which could impose tradeoffs on avoidance behaviour. The parasites’ distributions were highly contrasting, likely driven by different environmental factors: for example, sett microclimates favouring flea survival will facilitate their transmission [49,64]; similarly, *Eimeria* is transmitted faecal-orally, and oocysts are likely acquired from warm, moist sett chambers, which may explain the gradual decline moving away from the Thames river in the Northwest toward drier parts of the woods (Figure 1F). Ultimately, in combination with other socio-ecological factors (e.g., finding suitable mates), badgers may be incapable of completely avoiding all parasites via denning decisions, which could mediate co-infection and promote diverse parasite communities, giving rise to local hotspots across the population. This may lead to parasite avoidance being traded off against foraging, reproductive success and survival [22,71]. If badgers exhibit between-individual variation in movement and foraging specialisation [72], avoidance of multiple co-infecting parasites could maintain between-individual immune heterogeneity [55]. Notably, only cub density was negatively associated with lice. This observation could be linked to the detectable mortality effects in cubs, driving local cub mortality and/or motivating reproducing females to avoid spatial hotspots of lice.

All of the parasites we examined employ some degree of indirect transmission, likely yielding different relationships with host density than more directly transmitted parasites [1,17]. Ultimately, only avoiding conspecifics can help avoid direct parasite transmission (e.g. [8]), which may not be achievable through purely spatial structuring. Future studies could examine sexually transmitted infections or aerosol-transmitted viruses to investigate whether individuals living in areas of lower density gain any benefit in terms of direct parasite transmission. This may be of particular importance for bovine tuberculosis (bTB), which has a complex, nonlinear relationship with badger sociality [73–75]. Notably, previous bTB studies have generally used social group size as a metric of sociality; given that bTB has an environmental transmission component, particularly between badgers and cattle [76], spatial density metrics, such as those employed here, could be revealing.

We note two potential sources of negative density dependence untested by our modelling approach: the encounter-dilution effect [77,78] and micronutrient impacts on immunity [79]. For the former, where parasites exhibit a constant attack rate in space, greater host densities actually drive a lower per-individual burden of disease because the same number of parasites is divided among more available hosts [77,78]. Because this requires that the spatial distributions of parasites are not tightly linked to the host distribution (e.g. through reproducing on the host), it is unlikely to apply for any of the badger-specific parasites studied here. However, non-host-specific ticks (*Ixodes* sp.) could transmit from other species to badgers in a given area, producing an encounter-dilution effect; therefore, we are unable to rule out this mechanism (Table 2). Regarding nutritional status, we used body condition index as a coarse predictor of the ability to mount a healthy immune response. However, micronutrients are essential to effective immune function [79] and would not be detected from body condition indices; therefore, it is plausible that badgers might congregate in areas of high micronutrient availability. If this behavioural tendency functions to combat pathogens specifically, it amounts to a “landscape of disgust” acting through reduced susceptibility rather than reduced exposure. This possibility could be tested in this or similar systems by comparing the spatial distributions of high-resolution individual-level immune measures with host density distributions.

Our findings have important implications for the socio-spatial dynamics of this system and its resilience to pathogens and ecological change. If badger populations are organised optimally to occupy areas of least parasite transmission, even small disturbances (e.g. setts lost to forestry) could disrupt its socio-spatial structure and force individuals into unfavourable, more highly parasitised areas, exacerbating their disease burden. Therefore, disruptive anthropogenic activities such as culling-linked perturbation could have unseen consequences for badger disease beyond larger-scale movement impacts (e.g. [39,40]). These findings further inform our understanding of the drivers of badger spatial behaviour, offering insights that may be invaluable for their conservation and disease control [38,46].

## Acknowledgements

We would like to thank two anonymous reviewers, and Dan Becker, Amy Sweeny, and Quinn Webber for comments on the manuscript. We also warmly acknowledge the many students and co-workers who helped to collect this substantial data set. GFA and SB were supported by NSF grant number 1414296.

## Supplementary figures and tables: Negative density-dependent parasitism in a group-living carnivore

**Figure SI1:**
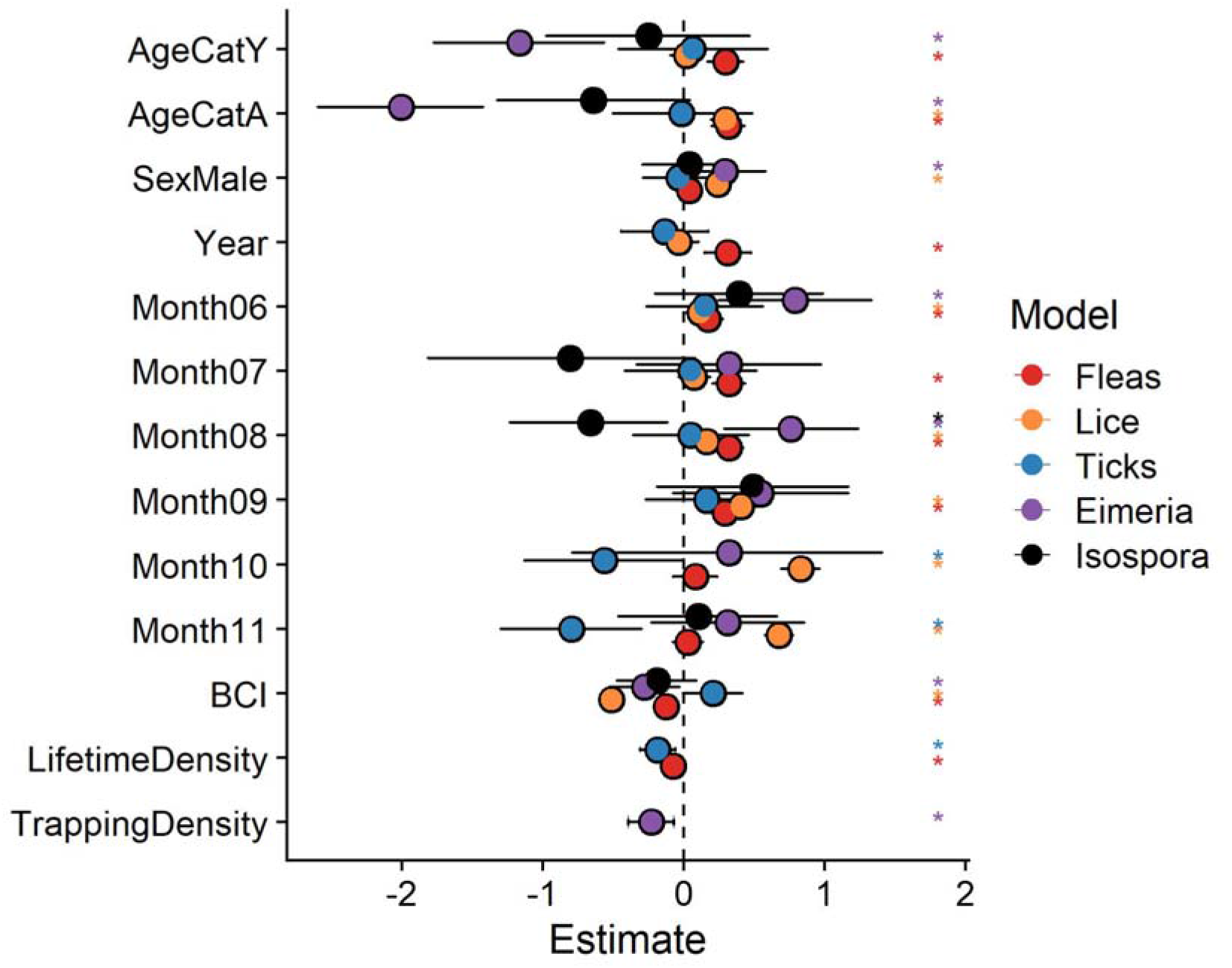
Model effect outputs for the full dataset INLA density models. Points represent modes of the posterior distribution of the effect estimates; error bars represent the 95% credibility intervals. Asterisks denote significant results, whose 95% credibility intervals did not overlap with zero. Only the behavioural traits that improved the model fit (ΔDIC<-) and were kept in the final model are shown.

**Figure SI2:**
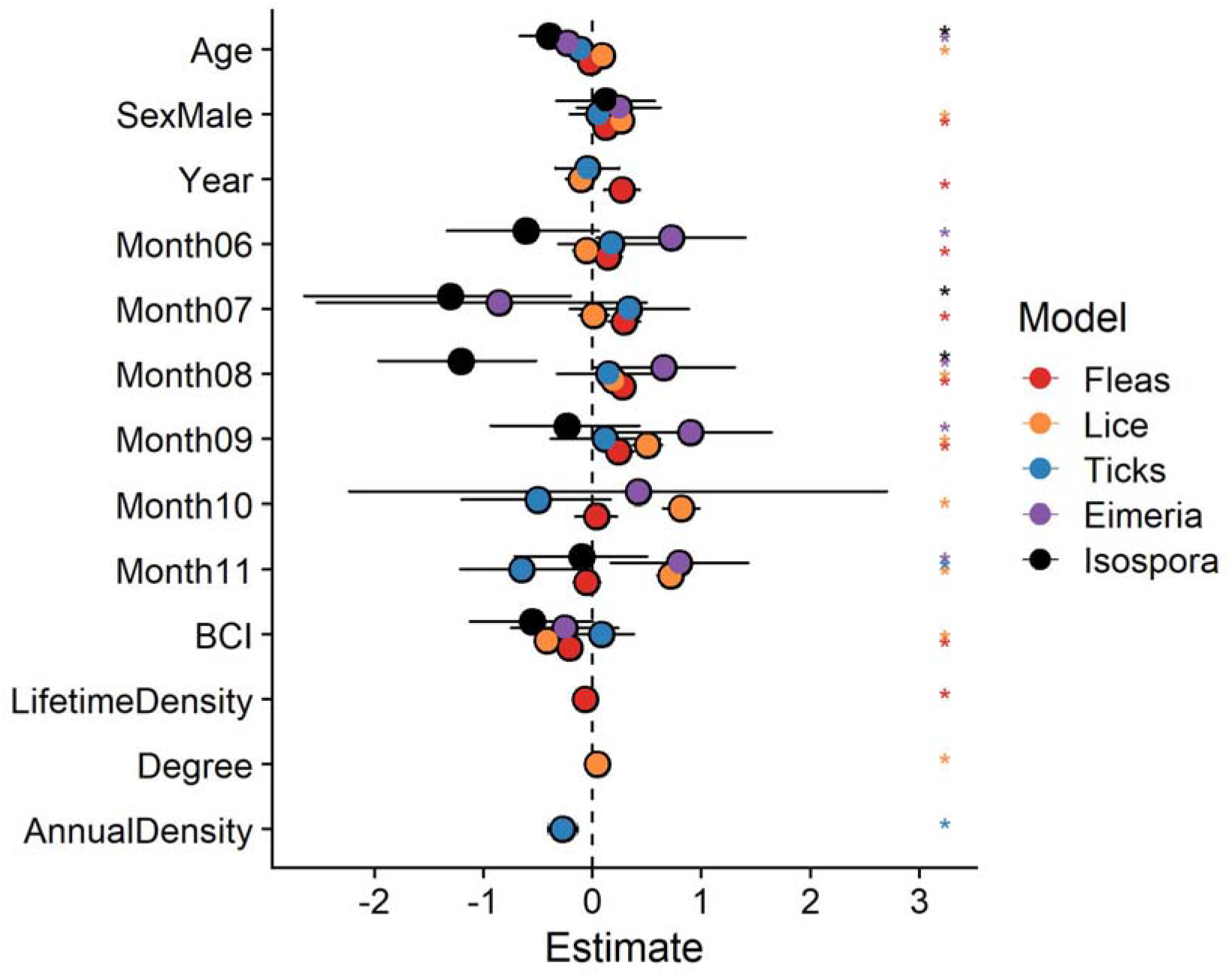
Model effect outputs for the adults-only dataset INLA density models. Points represent modes of the posterior distribution of the effect estimates; error bars represent the 95% credibility intervals. Asterisks denote significant results, whose 95% credibility intervals did not overlap with zero. Only the behavioural traits that improved the model fit (ΔDIC<-2) and were kept in the final model are shown.

**Figure SI3:**
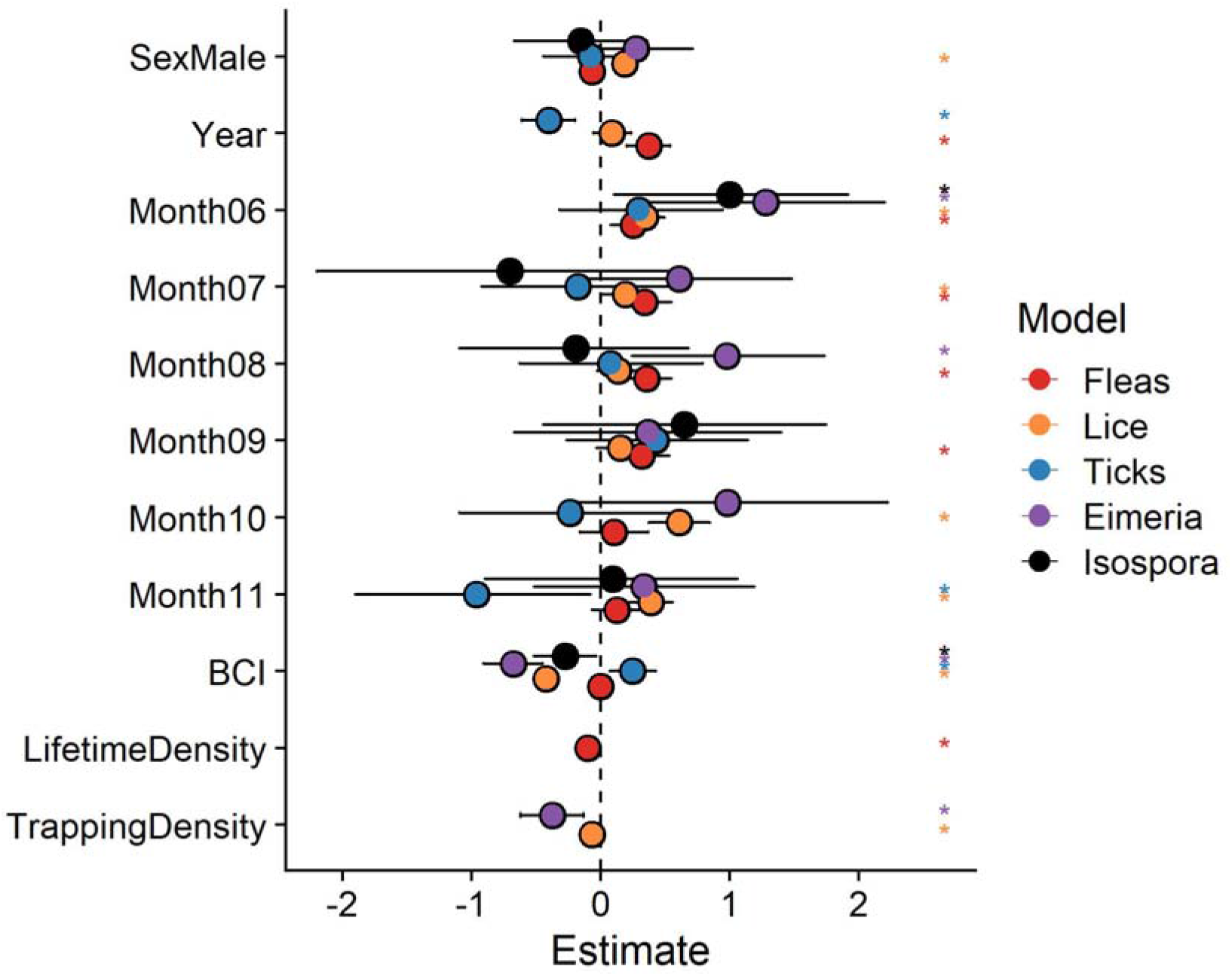
Model effect outputs for the juveniles-only dataset INLA density models. Points represent modes of the posterior distribution of the effect estimates; error bars represent the 95% credibility intervals. Asterisks denote significant results, whose 95% credibility intervals did not overlap with zero. Only the behavioural traits that improved the model fit (ΔDIC<-2) and were kept in the final model are shown.

**Figure SI4:**
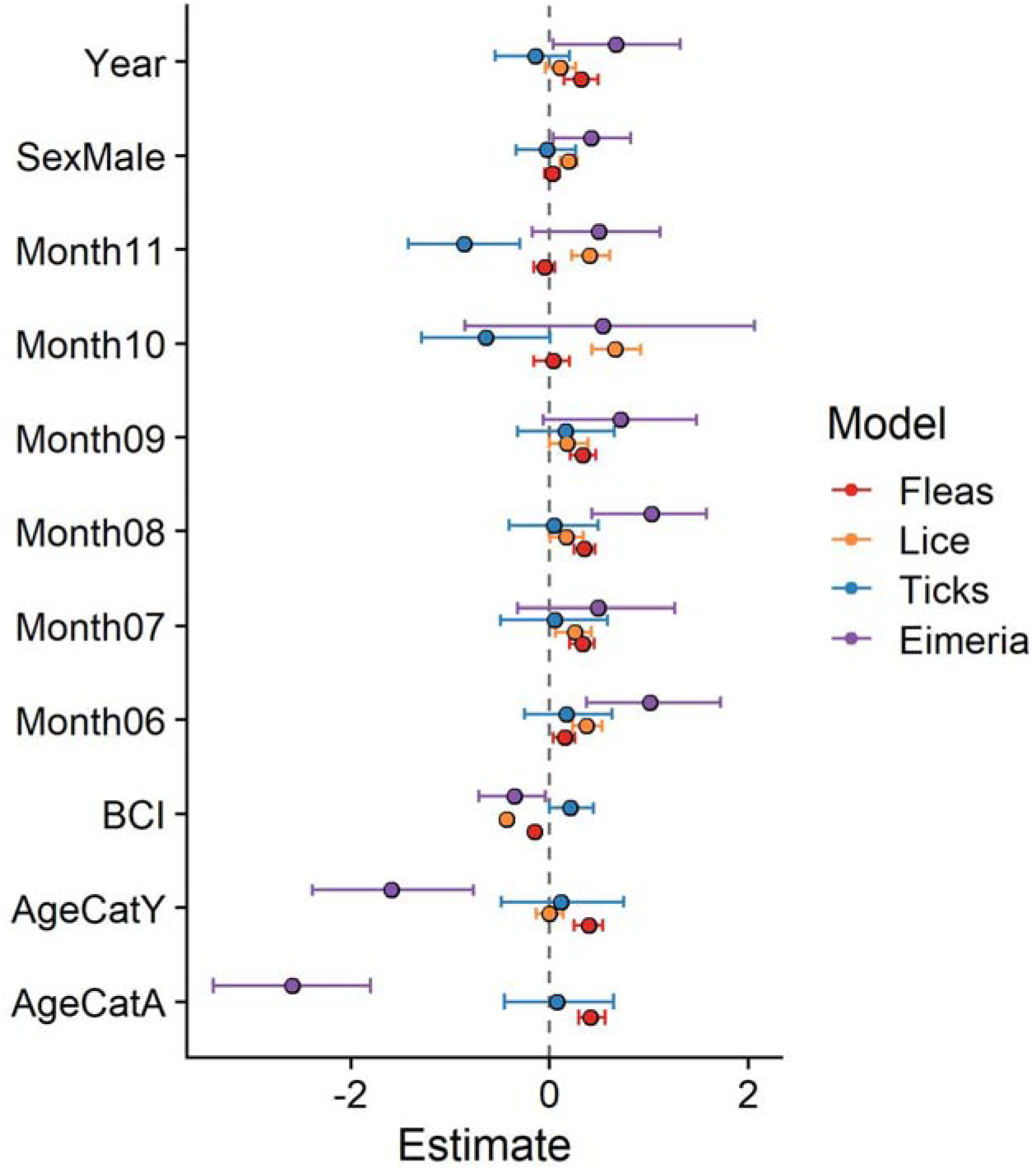
Model effect outputs for the multivariate MCMCglmm models closely represent the INLA univariate models. Points represent modes of the posterior distribution of the effect estimates; error bars represent the 95% credibility intervals. Asterisks denote significant results, whose 95% credibility intervals did not overlap with zero. Only the behavioural traits that improved the model fit (ΔDIC<-2) and were kept in the final model are shown.

**Table SI1:** DIC Changes (ΔDIC) associated with density measure model addition for the full models. Negative ΔDIC values correspnd to increased model fit. In each round, the variable that increased model the most was retained, and then the process was repeated, until no variables improved the model by decreasing ΔDIC by more than −2. Variables that were retained in each round are highlighted by asterisks.

| PARASITE | ROUND | VARIABLE | DELTA | KEPT |
| --- | --- | --- | --- | --- |
| FLEAS | 1 | Degree | 2.08 | 0 |
|  | 1 | GroupSize | 2.055 | 0 |
|  | 1 | **LifetimeDensity** | -16.001 | 1 |
|  | 1 | TrappingDensity | 1.115 | 0 |
|  | 1 | AnnualDensity | -6.889 | 0 |
| LICE | 1 | Degree | 1.796 | 0 |
|  | 1 | GroupSize | 1.3 | 0 |
|  | 1 | LifetimeDensity | 1.694 | 0 |
|  | 1 | TrappingDensity | 0.166 | 0 |
|  | 1 | AnnualDensity | 1.117 | 0 |
| TICKS | 1 | Degree | 0.773 | 0 |
|  | 1 | GroupSize | 3.644 | 0 |
|  | 1 | **LifetimeDensity** | -2.192 | 1 |
|  | 1 | TrappingDensity | 1.007 | 0 |
|  | 1 | AnnualDensity | -0.266 | 0 |
| EIMERIA | 1 | Degree | 1.134 | 0 |
|  | 1 | GroupSize | 1.79 | 0 |
|  | 1 | LifetimeDensity | 1.084 | 0 |
|  | 1 | **TrappingDensity** | -6.035 | 1 |
|  | 1 | AnnualDensity | 1.851 | 0 |
| ISOSPORA | 1 | Degree | 1.897 | 0 |
|  | 1 | GroupSize | 1.861 | 0 |
|  | 1 | LifetimeDensity | 1.873 | 0 |
|  | 1 | TrappingDensity | 1.507 | 0 |
|  | 1 | AnnualDensity | 1.717 | 0 |

**Table SI2:** DIC Changes (ΔDIC) associated with density measure model addition for the adults-only models. Negative ΔDIC values correspond to increased model fit. In each round, the variable that increased model the most was retained, and then the process was repeated, until no variables improved the model by decreasing ΔDIC by more than −2. Variables that were retained in each round are highlighted by asterisks.

| PARASITE | ROUND | VARIABLE | DELTA | KEPT |
| --- | --- | --- | --- | --- |
| FLEAS | 1 | Degree | 1.835 | 0 |
|  | 1 | GroupSize | 1.251 | 0 |
|  | 1 | **LifetimeDensity** | -8.444 | 1 |
|  | 1 | TrappingDensity | 1.509 | 0 |
|  | 1 | AnnualDensity | -4.549 | 0 |
| LICE | 1 | **Degree** | -2.388 | 1 |
|  | 1 | GroupSize | 1.747 | 0 |
|  | 1 | LifetimeDensity | 1.332 | 0 |
|  | 1 | TrappingDensity | 1.943 | 0 |
|  | 1 | AnnualDensity | 0.944 | 0 |
| TICKS | 1 | Degree | 1.93 | 0 |
|  | 1 | GroupSize | -12.573 | 0 |
|  | 1 | LifetimeDensity | -5.864 | 0 |
|  | 1 | TrappingDensity | 1.2 | 0 |
|  | 1 | **AnnualDensity** | -14.067 | 1 |
| EIMERIA | 1 | Degree | 14.117 | 0 |
|  | 1 | GroupSize | 1.964 | 0 |
|  | 1 | LifetimeDensity | 1.817 | 0 |
|  | 1 | TrappingDensity | -0.283 | 0 |
|  | 1 | AnnualDensity | 1.237 | 0 |
| ISOSPORA | 1 | Degree | 1.569 | 0 |
|  | 1 | GroupSize | 0.882 | 0 |
|  | 1 | LifetimeDensity | 1.9 | 0 |
|  | 1 | TrappingDensity | 1.65 | 0 |
|  | 1 | AnnualDensity | 1.8 | 0 |

**Table SI3:** DIC Changes (ΔDIC) associated with density measure model addition for the juvenile-only models. Negative ΔDIC values correspond to increased model fit. In each round, the variable that increased model the most was retained, and then the process was repeated, until no variables improved the model by decreasing ΔDIC by more than −2. Variables that were retained in each round are highlighted by asterisks.

| PARASITE | ROUND | VARIABLE | DELTA | KEPT |
| --- | --- | --- | --- | --- |
| FLEAS | 1 | Degree | 0.985 | 0 |
|  | 1 | GroupSize | -2.362 | 0 |
|  | 1 | **LifetimeDensity** | -15.209 | 1 |
|  | 1 | TrappingDensity | 1.337 | 0 |
|  | 1 | AnnualDensity | -10.016 | 0 |
| LICE | 1 | Degree | -2.862 | 0 |
|  | 1 | GroupSize | 1.747 | 0 |
|  | 1 | LifetimeDensity | -3.907 | 0 |
|  | 1 | **TrappingDensity** | -8.087 | 1 |
|  | 1 | AnnualDensity | -2.572 | 0 |
| TICKS | 1 | Degree | 1.602 | 0 |
|  | 1 | GroupSize | 0.462 | 0 |
|  | 1 | LifetimeDensity | -0.244 | 0 |
|  | 1 | TrappingDensity | 0.918 | 0 |
|  | 1 | AnnualDensity | -0.459 | 0 |
| EIMERIA | 1 | Degree | 1.42 | 0 |
|  | 1 | GroupSize | 1.067 | 0 |
|  | 1 | LifetimeDensity | -0.127 | 0 |
|  | 1 | **TrappingDensity** | -7.465 | 1 |
|  | 1 | AnnualDensity | -0.945 | 0 |
| ISOSPORA | 1 | Degree | 1.73 | 0 |
|  | 1 | GroupSize | 1.096 | 0 |
|  | 1 | LifetimeDensity | 1.775 | 0 |
|  | 1 | TrappingDensity | 1.567 | 0 |
|  | 1 | AnnualDensity | 1.709 | 0 |

